# Reexamination of N-terminal domains of Syntaxin-1 in vesicle fusion from central murine synapses

**DOI:** 10.1101/2021.02.15.431257

**Authors:** Gülçin Vardar, Andrea Salazar-Lázaro, Marisa Brockmann, Marion Weber-Boyvat, Sina Zobel, Victor Wumbor-Apin Kumbol, Thorsten Trimbuch, Christian Rosenmund

## Abstract

Syntaxin-1 (STX1) and Munc18-1 are two requisite components of synaptic vesicular release machinery, so much so synaptic transmission cannot proceed in their absence. They form a tight binary complex through two major binding modes: one through STX1’s N-peptide, the other through STX1’s closed conformation driven by its H_abc_-domain. However, physiological roles of these two reportedly different binding modes in synapses are still controversial. Here we characterized the roles of STX1’s N-peptide, H_abc_-domain, and open conformation with and without N-peptide deletion using our STX1-null mouse model system and exogenous reintroduction of STX1A mutants. We show, on the contrary to the general view, that the H_abc_-domain is absolutely required and N-peptide is dispensable for synaptic transmission. However, STX1’s N-peptide plays a regulatory role, particularly in the Ca^2+^-sensitivity and the short-term plasticity of vesicular release, whereas STX1’s open conformation governs the vesicle fusogenicity. Strikingly, we also show that neurotransmitter release still proceeds when both the interaction modes between STX1 and Munc18-1 are presumably intervened together, necessitating a refinement of the conceptualization of STX1–Munc18-1 interaction.

## Introduction

The synaptic vesicle (SV) fusion is the fundamental process in synaptic transmission and it is catalyzed by the merger of plasma and vesicular membranes by the neuronal SNAREs syntaxin-1 (STX1 collectively refers to STX1A and STX1B throughout this study), synaptobrevin-2 (Syb-2), and SNAP25 (Baker & Hughson, 2016; Rizo & Sudhof, 2012; Rizo & Xu, 2015). STX1 is the most important neuronal SNARE because synaptic transmission grinds to a halt in its absence (Vardar *et al*, 2016). Compared to the other SNAREs, it also has a unique structure with its regulatory region composed of a bulky three helical H_abc_-domain and a short N-peptide preceding its SNARE motif (Fig 1A) (Fernandez *et al*, 1998).

**Figure 1:**
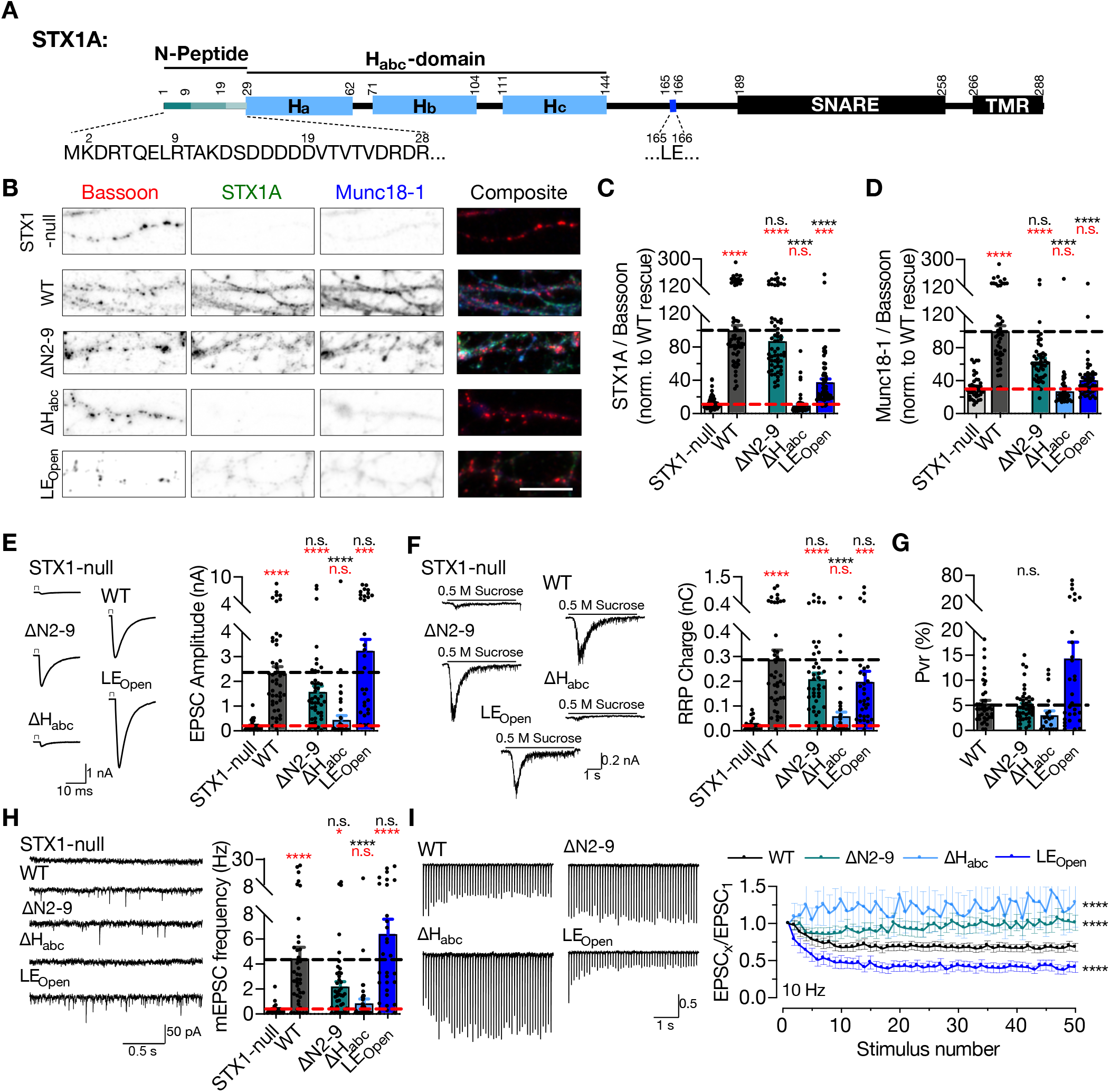
STX1’s H_abc_-domain is essential and N-peptide is dispensable for neurotransmitter release. A. Domain structure of STX1A. The protein consists of a short N-Peptide (aa 1-9 or 1-28), H_abc_ domain (aa 29-144) formed by three helices, H_a_, H_b_, and H_c_, followed by the H3 helix (aa 189 - 259; SNARE domain) and a transmembrane region (aa 266-288; TMR). B. Example images of immunofluorescence labeling for Bassoon, STX1A, and Munc18-1, shown as red, green, and blue, respectively, in the corresponding composite pseudocolored images obtained from high-density cultures of STX1-null hippocampal neurons either not rescued or rescued with STX1A^WT^, or STX1A^Δ2-9^; STX1A^LEOpen^; or STX1A^ΔHabc^. Scale bar: 10 μm C-D. Quantification of the immunofluorescence intensity of STX1A and Munc18-1 as normalized to the immunofluorescence intensity of Bassoon in the same ROIs as shown in (B). The values were then normalized to the values obtained from STX1A^WT^ neurons. E. Example traces (left) and quantification of the amplitude (right) of EPSCs obtained from hippocampal autaptic STX1-null neurons either not rescued or rescued with STX1A^WT^, STX1B^Δ2-9^, STX1A^LEOpen^, or STX1A^ΔHabc^. F. Example traces (left) and quantification of the charge transfer (right) of 500 mM sucrose-elicited RRPs obtained from the same neurons as in E. G. Quantification of Pvr determined as the percentage of the RRP released upon one AP. H. Example traces (left) and quantification of the frequency (right) of mEPSCs recorded at −70 mV. I. Example traces (left) and quantification (right) of STP determined by high-frequency stimulation at 10 Hz and normalized to the EPSC_1_ from the same neuron. Data information: The artifacts are blanked in example traces in (D) and (H). The example traces in (G) were filtered at 1 kHz. In (C–H), data points represent single observations, the bars represent the mean ± SEM. In (I), data points represent mean ± SEM. Red and black annotations (stars and n.s.) on the graphs show the significance comparisons to STX1-null and to STX1A^WT^ rescue, respectively (nonparametric Kruskal-Wallis test followed by Dunn’s *post hoc* test, *p ≤ 0.05, ***p ≤ 0.001, ****p ≤ 0.0001. Two-way ANOVA was applied for data in (I). The numerical values are summarized in source data.

Besides its interaction with the other SNAREs, STX1 also binds to its cognate SM protein Munc18-1 forming a tight binary complex with an affinity in the nanomolar range (Burkhardt *et al*, 2008; Pevsner *et al*, 1994). Munc18-1, which is an assistor of SNARE-mediated vesicular release, is an equally important protein as its absence also leads to inhibition of synaptic transmission (Verhage *et al*, 2000). Two major modes for STX1 binding to Munc18-1 have been defined: one through its N-peptide, the other through its closed conformation driven by the intramolecular interaction between its H_abc_- and SNARE domains (Dulubova *et al*, 1999; Misura *et al*, 2000). However, several issues regarding these reportedly different binding modes of STX1 to Munc18-1 are still subjects of dispute.

It is evident that STX1’s H_abc_-domain is required for proper folding of STX1 and for proper co-recruitment of STX1–Munc18-1 complex to the active zone (AZ) (Han *et al*, 2009; Meijer *et al*, 2012; Vardar *et al*, 2020; Zhou *et al*, 2013), yet it has been deemed to play a secondary role in synaptic transmission, to the point that it is dispensable for vesicle fusion *per se* (Meijer *et al.*, 2012; Rathore *et al*, 2010; Shen *et al*, 2010; Zhou *et al.*, 2013). However, an increasing number of mutations discovered in the H_abc_-domain of STX1B in patients with epilepsy (Schubert *et al*, 2014; Vardar *et al.*, 2020; Wolking *et al*, 2019) points to greater importance for this region in neurotransmitter release.

The physiological significance of Munc18-1 binding to STX1’s N-peptide is less clear, even though the general view leans towards its indispensability for synaptic transmission. Firstly, The STX1 N-peptide does not majorly contribute to its overall affinity for Munc18-1 (Burkhardt *et al.*, 2008; Christie *et al*, 2012; Colbert *et al*, 2013), yet liposome fusion cannot proceed without the N-peptide in reconstitution experiments (Rathore *et al.*, 2010; Shen *et al.*, 2010; Shen *et al*, 2007). On the other hand, interfering with STX1-N-peptide–Munc18-1 interaction by mutations either on STX1 (Zhou *et al.*, 2013) or on Munc18-1 (Han *et al.*, 2009; Khvotchev *et al*, 2007; Meijer *et al.*; Park *et al*, 2016; Shen *et al.*, 2007) in synapses in diverse model systems disclosed either its essentiality or its dispensability. Thus, a collective consensus as to what function the binding of STX1’s highly conserved N-peptide to Munc18-1 plays in synaptic transmission has not been reached.

So far, the physiological roles of STX1’s N-peptide, H_abc_-domain, and open-closed conformation were not assessed in central synapses completely devoid of STX1. Rather, studies have been conducted either in synapses with normal STX1 expression but mutant Munc18-1 (Khvotchev *et al.*, 2007; Meijer *et al.*, 2012; Shen *et al*, 2018) or in synapses with only severely reduced expression of STX1 (Zhou *et al.*, 2013). Furthermore, *in vitro* studies do not contain the full panel of native synaptic proteins and mostly do not use full-length STX1 (Rathore *et al.*, 2010; Shen *et al.*, 2010; Shen *et al.*, 2007). Therefore, we addressed the contribution of different domains of STX1 to neurotransmission using our STX1-null mouse model system and exogenous reintroduction of STX1A mutants either lacking N-peptide or the H_abc_-domain, or STX1 mutants forced into the open conformation (LE_Open_-mutation) with or without an N-peptide deletion. We show that the H_abc_-domain is absolutely required for STX1’s stability and/or expression and thus neurotransmitter release. Furthermore, in contrast to the general view, we find that N-peptide is not indispensable for synaptic transmission, however, propose that STX1’s N-peptide plays a regulatory role, particularly in the Ca^2+^-sensitivity of vesicular release and generally in vesicle fusion, which is only unmasked by STX1’s open conformation.

## Results

### STX1’s H_abc_-domain is essential and N-peptide is dispensable for neurotransmitter release

Vesicle fusion does not occur in the absence of STX1 (Vardar *et al.*, 2016) providing a null background in terms of neurotransmitter release. Thus, we used STX1A constitutive, STX1B conditional knockout (STX1-null) mouse neurons and lentiviral expression of different STX1 mutants in conjunction with *Cre* recombinase (Vardar *et al.*, 2016; Vardar *et al.*, 2020) to study the structure-function relationship of STX1 domains. With the focus on the effects of different Munc18-1 binding modes, we expressed STX1A mutants either with the deletion of the N-peptide (STX1A^ΔN2-9^) or the H_abc_-domain (Δ29-144; STX1A^ΔHabc^) or with the introduction of well-described LE_Open_ (L165A, E166A; STX1A^LEOpen^) mutation (Fig 1A).

Firstly, we utilized immunocytochemistry in high-density hippocampal neuronal culture to quantify the exogenous expression of STX1A^ΔN2-9^, STX1A^ΔHabc^, and STX1A^LEOpen^ at presynaptic compartments as defined by Bassoon-positive puncta and normalized fluorescence signals to the signals caused by expression of STX1A^WT^, all in STX1-null neurons. As expected from previous studies (Meijer *et al.*, 2012; Zhou *et al.*, 2013), deletion of the N-peptide had no significant effect on STX1A expression compared to STX1A^WT^, whereas STX1A^ΔHabc^ did not produce a measurable signal (Fig 1B & 1C). Loss of STX1 leads to a severe reduction in Munc18-1 expression, which can be rescued by the expression of either STX1A or STX1B (Vardar *et al.*, 2016; Vardar *et al.*, 2020; Zhou *et al.*, 2013). Consistent with the expected relative binding states of STX1A^ΔN2-9^ and STX1A^ΔHabc^ to Munc18-1, N-peptide deletion did not cause a significant change in Munc18-1 expression at Bassoon positive puncta whereas the H_abc_-domain deletion was unable to rescue Munc18-1 levels back to WT-like levels (Fig 1B & 1D). Rendering STX1B constitutively open by LE_Open_-mutation is also known to decrease STX1B as well as Munc18-1 levels (Gerber *et al*, 2008) and the expression of STX1A^LEOpen^ was severely low and inefficient to rescue Munc18-1 levels (Fig 1B, 1C, &1D).

To assess how the interruption of the different Munc18-1 binding domains of STX1A affect the release of presynaptic vesicles, we measured Ca^2+^-triggered and spontaneous vesicle fusion, vesicle priming, and short-term plasticity in autaptic hippocampal neurons using electrophysiology as described previously (Vardar *et al.*, 2016; Vardar *et al.*, 2020). Consistent with the previous studies on the constitutively ‘open’ STX1A or STX1B (Gerber *et al.*, 2008; Zhou *et al.*, 2013), the EPSCs trended towards an increase (Fig 1E), and the RRP assessed by hypertonic sucrose application (Fig 1F) trended towards a decrease (Fig. 1F) leading to a significant increase in Pvr (Fig 1G). The increase in Pvr was also evident in the observed enhancement of short-term depression (Fig 1I) as well as in the trend toward increased mEPSC frequency (Fig 1H). These findings support the robustness of our model-system.

Surprisingly, loss of N-peptide of STX1A did not produce any significant change in Ca^2+^-evoked vesicular release, as the EPSCs were comparable between STX1A^WT^ and STX1A^ΔN2-9^ neurons (Fig 1E), on the contrary to the previous studies (Zhou *et al.*, 2013). We also did not detect any significant alterations in the size of the RRP nor in the spontaneous neurotransmission upon deletion of STX1’s N-peptide (Fig 1E & 1F), but only a trend towards a decrease. Proportionally similar trends in the reduction of both EPSC and RRP resulted in comparable Pvr between STX1A^ΔN2-9^ and STX1A^WT^ neurons (Fig 1G). Despite the lack of a net difference in the Pvr, however, STX1A^ΔN2-9^ neurons exhibited an altered short-term plasticity (STP) in response to the 10 Hz stimulation, with no depression to latter stimuli (Fig 1I).

Previous studies have suggested that the H_abc_-domain of STX1A and particularly its interaction with Munc18-1 is dispensable for vesicle fusion both *in vitro* and *in vivo* (Meijer *et al.*, 2012; Rathore *et al.*, 2010; Shen *et al.*, 2010; Zhou *et al.*, 2013). However, our analysis of the neurotransmission properties of the STX1A^ΔHabc^ neurons in comparison to the STX1A^WT^ neurons showed that STX1A^ΔHabc^ was incapable of rescuing neurotransmitter release as it produced no detectable EPSC, RRP, or mEPSC; a phenotype similar to the STX1-null neurons (Fig 1E, 1F, & 1G).

### STX1 H_abc_-domain is indispensable for neuronal viability and the organization of synaptic ultrastructure

STX1 has also an obligatory function in neuronal maintenance and complete loss of both STX1A and STX1B leads to neuronal death (Vardar *et al.*, 2016). To address the overall functionality of STX1A, we assessed the survivability of the high-density cultured STX1-null neurons expressing STX1A^ΔHabc^ and determined the cell number at different time intervals starting at DIV 8 (Fig 2A-2C), at which time point all the groups had an average of ~40 neurons per mm^2^ (Fig 2B). Then we calculated the ratio of the cell number at DIV 15, 22, and 29 to the cell number at DIV 8 as a read-out for neuronal viability. STX1-null neurons showed a dramatic loss between DIV 8 and DIV 15 (Fig 2C) as reported before (Vardar *et al.*, 2016). Even though at DIV 15 the number of surviving STX1A^ΔHabc^ neurons were slightly but significantly higher compared to that in STX1-null group, eventually STX1A^ΔHabc^ failed to rescue neuronal survival as by DIV 22 almost all STX1A^ΔHabc^ neurons were dead (Fig 2C).

**Figure 2:**
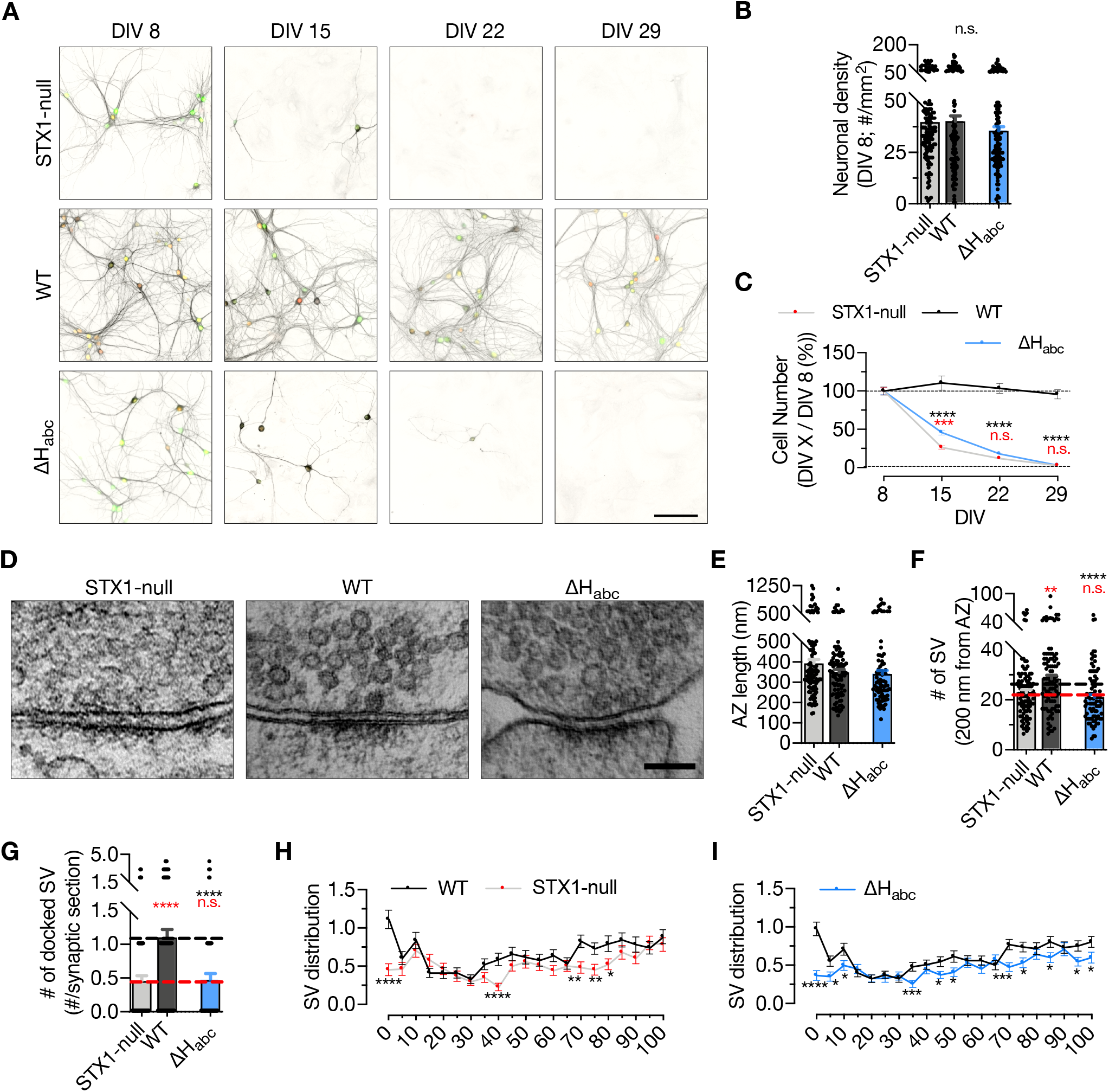
STX1’s H_abc_-domain is essential for the overall function of STX1. A. Example images of high-density cultures of STX1-null, STX1A^WT^, and STX1A^ΔHabc^ hippocampal neurons at DIV 8, 15, 22, and 29 represented with immunofluorescent labeling of MAP2. Red and green nuclei serve as a marker for NLS-RFP-P2A-*Cre* recombinase expression and for NLS-GFP-P2A-STX1A (either WT or mutants), respectively. Scale bar: 50 μm. B. Quantification of neuronal density at DIV 8. C. Quantification of the percentage of the surviving neurons at DIV 8, 15, 22, and 29 as normalized to the neuronal density at DIV 8 in the same well. D. Example HPF-EM images of nerve terminals from high-density cultures of STX1-null hippocampal neurons either not rescued or rescued with STX1A^WT^ or STX1A^ΔHabc^. E-F-G. Quantification of AZ length, number of SVs within 200nm distance from AZ, and number of docked SVs. H-I. SV distribution of STX1-null and STX1A^ΔHabc^ neurons compared to that of STX1A^WT^ neurons. Data information: In (B & E - G), data points represent single observations, the bars represent the mean ± SEM. In (C, H, & I), data points represent the mean ± SEM. Red and black annotations (stars and n.s.) on the graphs show the significance comparisons to STX1-null and to STX1A^WT^ neurons, respectively (nonparametric Kruskal-Wallis test followed by Dunn’s *post hoc* test, *p ≤ 0.05, **p ≤ 0.01, ***p ≤ 0.001, ****p ≤ 0.0001). The numerical values are summarized in source data.

Furthermore, we analyzed vesicle docking by morphological assessment of synaptic ultrastructure to determine whether STX1A^ΔHabc^ expression could reverse the impairment in the vesicle docking observed in STX1-null neurons (Vardar *et al.*, 2016). To circumvent the reduction in cell number and the synapse number thereof, we transduced the neurons at DIV 2-3 to postpone the cell death as previously shown (Vardar *et al.*, 2016), and analyzed the synapses using high-pressure freezing fixation (DIV 14-16) combined with electron microscopy (HPF-EM; Fig 2D-2H). Firstly, we observed no difference in the AZ length among STX1-null, STX1A^WT^, and STX1A^ΔHabc^ neurons (Fig 2E). On the other hand, the total SV number within 200 nm from the AZ was significantly reduced in STX1A^ΔHabc^ neurons compared to that in STX1A^WT^ neurons (Fig 2F). STX1A^ΔHabc^ also did not restore vesicle docking, which remained at ~50% of the STX1A^WT^ neurons (Fig 2G). Similarly, the SV distribution within 100 nm of the AZ were comparable between STX1-null and STX1A^ΔHabc^ neurons, with both significantly altered number of SVs compared to the STX1^WT^ neurons, especially in the 15, 40, and 100 nm range from AZ (Fig 2G & 2H). This suggests a general alteration of the synaptic organization even though the length of AZs were unaltered.

Based on the lack of immunofluorescent signal (Fig 1C) together with the lack of any rescue activity in any release parameters (Fig. 1E-1I, 2G & 2I) and neuronal survivability for STX1A^ΔHabc^ (Fig 2C), we again examined the expression level of STX1A^ΔHabc^ in comparison with STX1A^WT^, this time using constructs with a C-terminal FLAG tag (Fig EV1). C-terminal FLAG tag did not reveal significant changes in the expression of STX1A^WT^ (Fig EV1). We then measured the immunofluorescent signal using a FLAG antibody in the neurons expressing FLAG-tagged STX1A^WT^, STX1A^ΔN2-9^, STX1A^LEOpen^, or STX1A^ΔHabc^, all of which showed similar levels of reduction in the expression as compared to the non-tagged constructs (Fig EV1) suggesting that the lack of immunofluorescent signal in STX1A^ΔHabc^ (Fig 1C) is not due to a loss of antibody binding epitope, but rather due to the low level of protein.

### Deletion of the entire N-terminal stretch does not impair neurotransmitter release

It is striking that deletion of the 2-9 amino acids (aa), namely the N-peptide, of STX1A revealed no significant phenotype in synaptic transmission from central synapses (Fig. 1), even though this domain has been designated as a crucial factor for neurotransmitter release. Though the aa 2-9 has been defined as the residues binding to the outer surface of Munc18-1 (Burkhardt *et al.*, 2008; Hu *et al*, 2007), the whole 2-28 aa stretch manifests an unstructured nature in NMR studies (Misura *et al.*, 2000), suggesting a potential involvement in protein-protein interactions. Thus, we extended our analysis of the function of N-peptide by constructing STX1A with longer deletions in the N-terminus (STX1A^ΔN2-19^ and STX1A^ΔN2-28^), and probed the effects of these mutants on synaptic transmission.

Compared to the exogenous expression of STX1A^WT^, deletion of 19 or 28 aa from the N-terminus reduced the expression of STX1A to ~60% (Fig 3A-3B), suggesting a modulatory effect of the unstructured N-terminal domain on STX1’s expression or stability. However, neither the reduction in STX1A expression nor loss of the putative Munc18-1 binding domain influenced the Munc18-1 levels, which was effectively rescued back to WT-like levels (Fig 3A & 3C).

**Figure 3:**
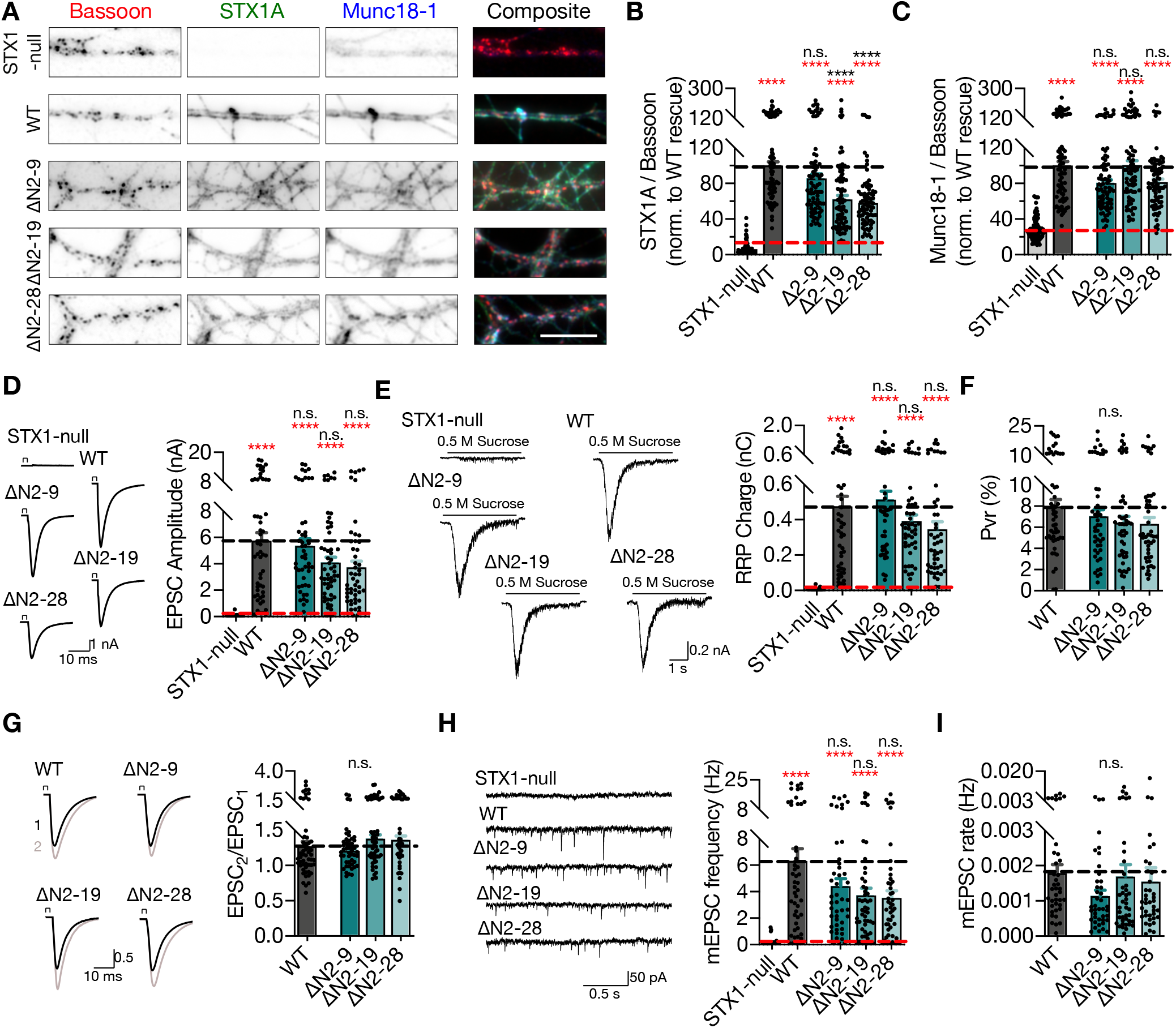
Deletion of the entire N-terminal stretch does not impair neurotransmitter release. A. Example images of immunofluorescence labeling for Bassoon, STX1A, and Munc18-1, shown as red, green, and blue, respectively, in the corresponding composite pseudocolored images obtained from high-density cultures of STX1-null hippocampal neurons either not rescued or rescued with STX1A^WT^, STX1A^Δ2-9^, STX1A^Δ2-19^, or STX1A^Δ2-28^. Scale bar: 10 μm B-C. Quantification of the immunofluorescence intensity of STX1A and Munc18-1 as normalized to the immunofluorescence intensity of Bassoon in the same ROIs as shown in (A). The values were then normalized to the values obtained from STX1A^WT^ neurons. D. Example traces (left) and quantification of the amplitude (right) of EPSCs obtained from hippocampal autaptic STX1-null neurons either not rescued or rescued with STX1A^WT^, STX1A^Δ2-9^, STX1A^Δ2-19^, or STX1A^Δ2-28^. E. Example traces (left) and quantification of the charge transfer (right) of sucrose-elicited RRPs obtained from the same neurons as in D. F. Quantification of Pvr determined as the percentage of the RRP released upon one AP. G. Example traces (left) and quantification (right) of PPR measured at 40 Hz. The artifacts are blanked in the example traces. H. Example traces (left) and quantification of the frequency (right) of mEPSCs. The example traces were filtered at 1 kHz. I. Quantification of mEPSC rate as spontaneous release of one unit of RRP. Data information: The artifacts are blanked in example traces in (D) and (G). The example traces in (H) were filtered at 1 kHz. In (B–I), data points represent single observations, the bars represent the mean ± SEM. Red and black annotations (stars and n.s.) on the graphs show the significance comparisons to STX1-null and to STX1A^WT^ neurons, respectively (nonparametric Kruskal-Wallis test followed by Dunn’s *post hoc* test, ****p ≤ 0.0001). The numerical values are summarized in source data.

Strikingly, similar to the deletion of the N-peptide, neither deletion of 2-19 aa nor 2-28 aa produced any significant effect on vesicle fusion nor on vesicle priming, only a slight but graded trend towards a decrease (Fig 3D-3H). STX1A^ΔN2-9^, STX1A^ΔN2-19^ and STX1A^ΔN2-28^ restored the Ca^2+^-triggered release and the size of the RRP back to around 4 nA and 0.4 nC, respectively, but showed a gradual trend towards a decrease compared to that of STX1A^WT^, which was around 6 nA and 0.5 nC. We observed no difference in spontaneous vesicle fusion rate, as determined by mEPSC frequencies, between STX1A^WT^ and STX1A^ΔN^ groups (Fig 3H).

A trend towards a reduction in release was also expressed in Pvr, such that STX1A^ΔN2-19^ and STX1A^ΔN2-28^ neurons manifested Pvr of ~6% whereas STX1A^WT^ and STX1A^ΔN2-9^ neurons released with a Pvr of ~8% and ~7%, respectively (Fig 3F). Similarly, spontaneous release inclined to be impaired but not significantly, remaining at around 4 Hz compared to 6 Hz of STX1A^WT^ (Fig 3G). As another measure of Pvr, we induced paired action potentials (APs) at 40 Hz and observed no difference in paired-pulse ratio (PPR) of EPSCs between STX1A^WT^ and STX1A^ΔN^ neurons (Fig 3I).

Apart from STX1A’s first 9 aa, the STX1-N-peptide—Munc18-1 interaction is also proposed to be regulated by the phosphorylation of STX1’s S14 residue by CKII (Rickman & Duncan, 2010). To test whether the phosphorylation of S14 affects Munc18-1 trafficking and neurotransmitter release, we generated phosphonull (S14A) and phosphomimetic (S14E) STX1A mutants. We again measured the STX1A and Munc18-1 levels at synapses, which revealed no impact of the phosphorylation status of S14 on either STX1A or Munc18-1 levels (Appendix Fig S1), consistent with the finding that S14A causes only a minor decrease in the affinity of STX1A to Munc18-1 (Burkhardt *et al.*, 2008). As a direct function of STX1A S14 phosphorylation on vesicular release from neurons or neuroendocrine cells has been also proposed (Rickman & Duncan, 2010; Shi *et al*, 2020), we tested whether it would also influence the fusion of presynaptic vesicles. Both STX1A^S14A^ and STX1A^S14E^ efficiently restored all the release parameters to WT-like levels in STX1-null neurons (Appendix Fig S1), which suggests that the modulation of the STX1A N-peptide–Munc18-1 interaction by S14 phosphorylation does not alter its function in neurotransmitter release from central synapses. Interestingly, STX1A^S14E^ led to a slight but significant increase in the degree of short-term depression compared to that of STX1A^WT^ (Appendix Fig S1). Neither N-peptide deletion nor phosphorylation modulation mutants compromised the neuronal survival (Appendix Fig S2).

### Interruption of both Munc18-1 binding modes of STX1 does not impair neurotransmitter release

Munc18-1 binding to the N-peptide or to the closed conformation of STX1 constitutes the two major interaction modes between these proteins, yet, neither mutation causes a deficit in synaptic release (Fig 1 & Fig 3). Even though Munc18-1 interacts with STX1A^WT^ through multiple interaction points including the SNARE motif of STX1A (Burkhardt *et al.*, 2008; Liang *et al*, 2013; Misura *et al.*, 2000), it is expected that the inhibition of both ‘closed’ and ‘N-peptide’ binding modes would result in a drastic loss of the STX1A–Munc18-1 binary complex (Rickman *et al*, 2007) and thereby a loss of neurotransmitter release. Thus, we constructed STX1A mutants in which the N-peptide is deleted at differing lengths in conjunction with the LE_Open_-mutation and firstly observed N-peptide deletion in addition to the LE_open_-mutation decreased the STX1A and Munc18-1 levels further than that already caused by LE_Open_-mutation alone (Fig 4A-4C).

**Figure 4:**
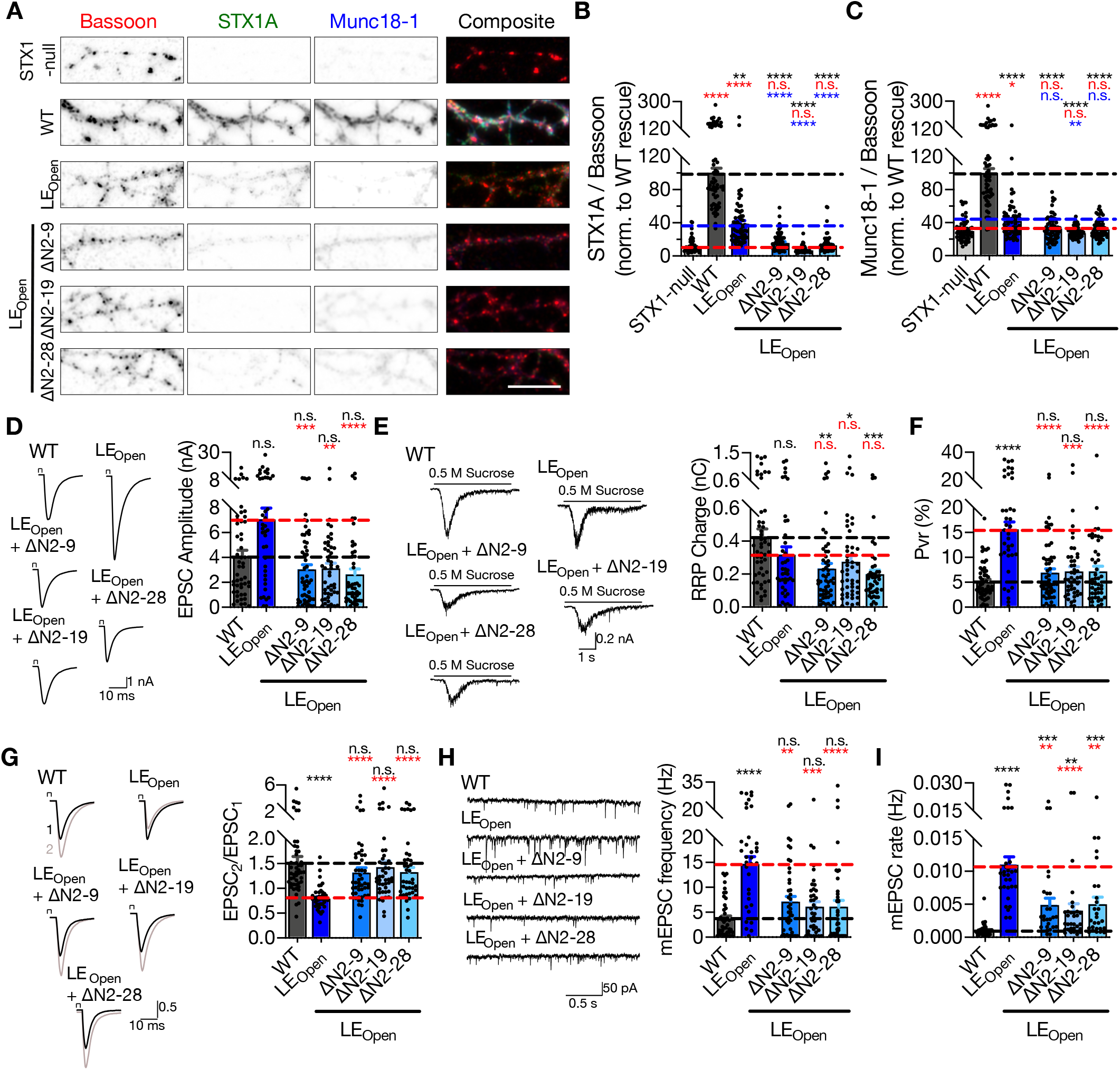
Interruption of both Munc18-1 binding modes of STX1 does not impair neurotransmitter release. A. Example images of immunofluorescence labeling for Bassoon, STX1A, and Munc18-1, shown as red, green, and blue, respectively, in the corresponding composite pseudocolored images obtained from high-density cultures of STX1-null hippocampal neurons either not rescued or rescued with STX1A^WT^, STX1A^LEOpen^, STX1A^LEOpen + ΔN2-9^, STX1A^LEOpen + ΔN2-19^, or STX1A^LEOpen + ΔN2-28^. Scale bar: 10 μm B-C. Quantification of the immunofluorescence intensity of STX1A and Munc18-1 as normalized to the immunofluorescence intensity of Bassoon in the same ROIs as shown in (A). The values were then normalized to the values obtained from STX1A^WT^ neurons. D. Example traces (left) and quantification of the amplitude (right) of EPSCs obtained from hippocampal autaptic STX1A^WT^, STX1A^LEOpen^, STX1A^LEOpen + ΔN2-9^, STX1A^LEOpen + ΔN2-19^, or STX1A^LEOpen + ΔN2-28^ neurons. E. Example traces (left) and quantification of the charge transfer (right) of sucrose-elicited RRPs obtained from the same neurons as in D. F. Quantification of Pvr determined as the percentage of the RRP released upon one AP. G. Example traces (left) and quantification (right) of PPR measured at 40 Hz. H. Example traces (left) and quantification of the frequency (right) of mEPSCs. I. Quantification of mEPSC rate as spontaneous release of one unit of RRP. Data information: The artifacts are blanked in example traces in (D) and (G). The example traces in (H) were filtered at 1 kHz. In (B–I), data points represent single observations, the bars represent the mean ± SEM. In (B & C), Red, black, and blue annotations (stars and n.s.) on the graphs show the significance comparisons to STX1-null, STX1A^WT^, and to STX1A^LEOpen^ neurons, respectively. In (D-I), red and black annotations on the graphs show the significance comparisons to STX1A^WT^, and STX1A^LEOpen^ neurons, respectively (nonparametric Kruskal-Wallis test followed by Dunn’s *post hoc* test, *p ≤ 0.05, **p ≤ 0.01, ***p ≤ 0.001, ****p ≤ 0.0001). The numerical values are summarized in source data.

Despite the presumed loss of STX1A–Munc18-1 interaction, it is remarkable that all LE_Open_-ΔN combination mutants rescued Ca^2+^-evoked neurotransmitter release to almost STX1A^WT^ levels with only a trend towards a reduction (Fig 4D). Because STX1A^LEOpen^ neurons showed increased EPSCs – albeit not significant – with an average of ~7 nA compared to ~4 nA of STX1A^WT^, EPSCs recorded from STX1A^LEOpen+ΔN^ neurons were significantly smaller than that of STX1A^LEOpen^ neurons and remained at ~3 nA (Fig 4D). This suggests that the small enhancement of Ca^2+^-evoked release by the presumed open conformation by LE_Open_-mutation was reversed by additional N-peptide deletions (Fig 4D). On the other hand, the reduction in RRP observed in neurons which express LE_Open_-mutation was not reverted back to WT-like levels by the addition of N-peptide deletions, but instead was further exaggerated as the RRP size significantly decreased in STX1A^LEOpen+ΔN^ neurons compared to that in STX1A^WT^ neurons (Fig 4E). As a result, increased Pvr, which is the hallmark phenotype of the LE_Open_-mutation (Gerber *et al.*, 2008), was reversed back to WT-like levels with only a trend toward a small increase (Fig 4F). This was also reflected in the decreased mEPSC frequency and in mEPSC release rate obtained from the STX1A^LEOpen+ΔN^ mutants compared to that of STX1A^LEOpen^ mutant (Fig 4G & 4H). Similarly, increased Pvr in STX1A^LEOpen^ expressing neurons led to decreased PPR when measured at 40 Hz, as N-peptide deletions in STX1A^LEOpen^ reverted PPR back to levels comparable to neurons expressing STX1A^WT^ (Fig 4I).

Surprisingly, putative disruption of the two supposedly main interaction points between STX1A and Munc18-1—by deleting N-peptide in its entirety in constitutively open STX1A—ultimately led to neuronal death (Fig EV2) indicative of independence of STX1’s functions in neurotransmitter release and neuronal maintenance of one another. However, the onset of cell death was postponed by the expression of STX1A^LEOpen+ΔN^ mutants compared to that observed in STX1-null neurons, as at DIV15 almost all neurons expressing the STX1A^LEOpen+ΔN^ mutants were still alive (Fig EV2). Because the electrophysiological recordings are mostly conducted at DIV 13 - 20, the compromised cell viability is unlikely to account for the reduction in neurotransmission in STX1A^LEOpen+ΔN^ mutants compared to that of in STX1A^LEOpen^ mutant.

A severe reduction in STX1 expression induced by *in vitro* knock-down (Arancillo *et al*, 2013; Zhou *et al.*, 2013) or transgenic knock-in (Arancillo *et al.*, 2013) strategies, results in a strong impairment in neurotransmitter release. Based upon that, we argued that the reduction of release parameters (Fig 4) of STX1A^LEOpen^ by additional N-peptide deletions may be due to decreased expression of STX1A (Fig 4B). To test this hypothesis, we down-titrated the viral load from 1X to 1/12X for STX1A^WT^ and to 1/3X and 1/6X for STX1A^LEOpen^ (Fig EV3). As our viral constructs include NLS-GFP before the P2A sequence followed by STX1A, nuclear GFP showed a decrease when the amount of virus was reduced (Fig EV3). Immunofluorescent labelling in autaptic neurons revealed that reducing the viral amount was effective in reducing expression levels of either STX1A^WT^ or STX1A^LEOpen^ to the levels comparable to that of STX1A^LEOpen+ΔN2-28^. However, reducing the expression level of STX1A^WT^ or STX1A^LEOpen^ did not cause a difference in their neurotransmitter release properties (Fig EV3).

It has been reported that Munc18-1 also functions upstream of the vesicle docking step (Gulyas-Kovacs *et al*, 2007; Toonen *et al*, 2006). Therefore, we analyzed the state of docked vesicles in neurons which express either STX1A^ΔN2-28^, STX1A^LEOpen^, or STX1A^LEOpen+ΔN2-28^ using HPF-EM (Fig 5A). AZ length was again comparable between all the mutants and STX1A^WT^ (Fig 5B). Strikingly, the neurons in which both Munc18-1 binding modes were inhibited by the STX1A^LEOpen+ΔN2-28^ mutation showed docked vesicles were reduced to ~50% of those in STX1A^WT^ synapses (Fig 5C). On the other hand, manipulation of either closed binding or N-peptide binding alone did not influence vesicle docking (Fig 5C). Previously, we have shown that vesicle priming can be completely abolished by a STX1A mutant (A240V, V244A) with the vesicle docking remaining intact (Vardar *et al.*, 2016). Furthermore, we also have reported that the vesicle priming is more prone to impairments by mutations in the vesicle release machinery than is vesicle docking, which suggests a separation or a different cooperativity between these events (Zarebidaki *et al*, 2020). In this light, we plotted the number of docked SVs versus the RRP size, and observed that the RRP is also more susceptible to a reduction than is vesicle docking for the STX1A– Munc18-1 binding mutants (Fig 5D). Vesicle distribution analysis revealed an accumulation of vesicles at 5nm distance from AZ in STX1A^ΔN2-28^ neurons but a reduction in STX1A^LEOpen+ΔN2-28^ neurons (Fig 5E & 5G), whereas STX1^LEOpen^ neurons did not show a major alteration in their vesicle distribution within 100nm from AZ (Fig 5F).

**Figure 5:**
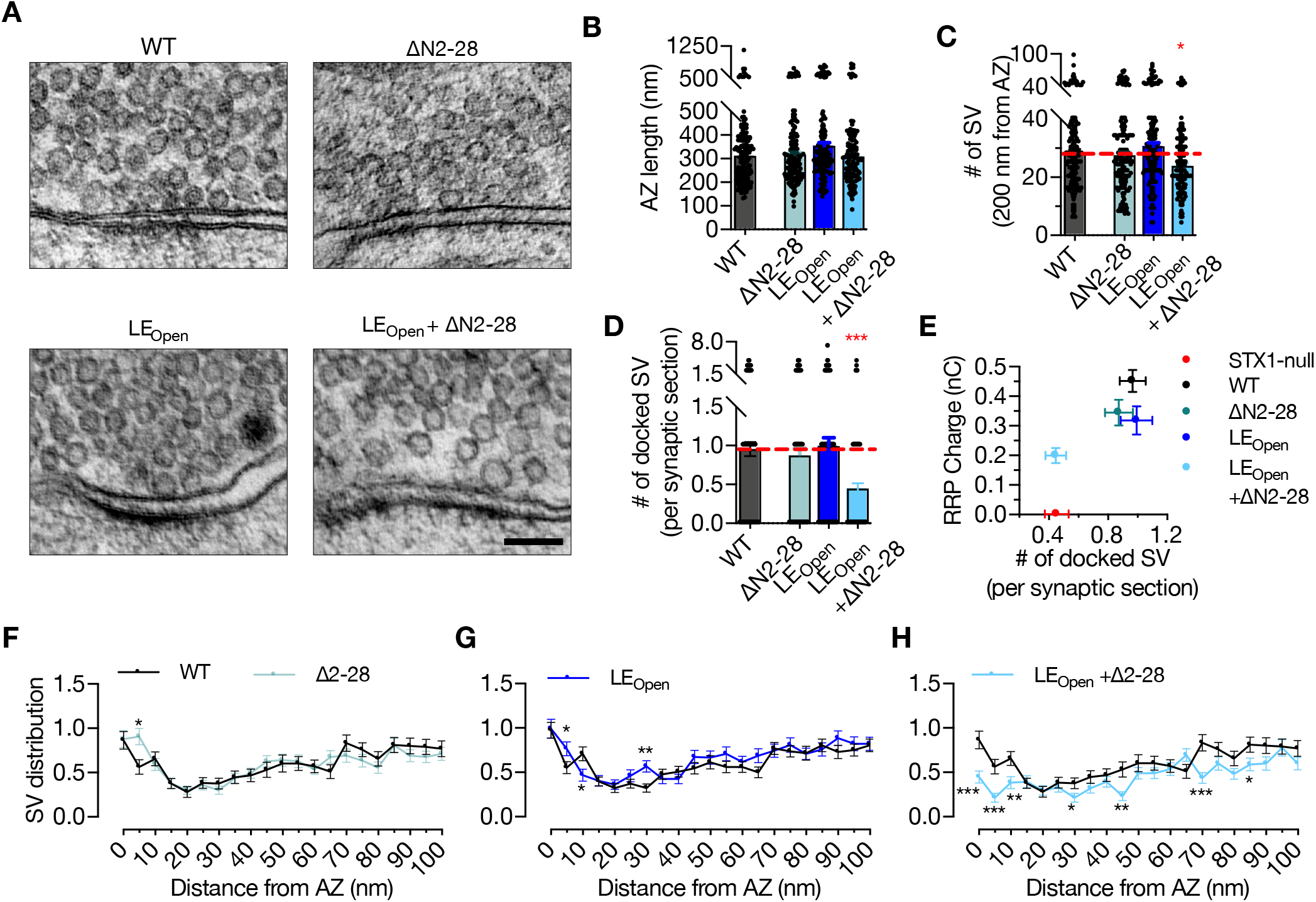
Interruption of both Munc18-1 binding modes of STX1 reduces the number of docked SVs. A. Example HPF-EM images of nerve terminals from high-density cultures of STX1A^WT^, STX1A^ΔN2-28^, STX1A^LEOpen^, and STX1A^LEOpen + ΔN2-28^ Neurons. B-C-D. Quantification of AZ length, number of SVs within 200nm distance from AZ, and number of docked SVs. E. Correlation of the number of docked SVs obtained by HPF-EM to the size of RRP obtained by electrophysiological recordings. F-G-H. SV distribution of STX1A^ΔN2-28^, STX1A^LEOpen^, and STX1A^LEOpen^ ^+ ΔN2-28^ neurons compared to that of STX1A^WT^ neurons. Data information: In (B–D), data points represent single observations, the bars represent the mean ± SEM. In (E-H), data points represent mean ± SEM. Black annotations on the graphs show the significance comparisons to STX1A^WT^ rescue (nonparametric Kruskal-Wallis test followed by Dunn’s *post hoc* test in (B-D), multiple t-tests in F-H *p ≤ 0.05, **p ≤ 0.01, ***p ≤ 0.001). The numerical values are summarized in source data.

### STX1’s N-peptide has a modulatory function in short-term plasticity and Ca^2+^-sensitivity of synaptic transmission

So far, our analysis has shown that STX1’s N-peptide is not indispensable for neurotransmitter release (Fig 1 and Fig 3), but plays a modulatory role in protein expression (Fig 3) and, when STX1 is in the open confirmation, in vesicle fusion and Pvr (Fig 4). To elucidate the modulation of neurotransmitter release by STX1’s N-peptide, we took a closer look at Pvr and its effect on STP (Fig 6A). Even though the neurons expressing any STX1A^ΔN^ mutant showed only a trend towards decreased Pvr compared to that of STX1A^WT^ neurons (Fig 4), their STP behavior in response to 50 stimuli at 10 Hz differed significantly. Both STX1A^ΔN2-19^ and STX1A^ΔN2-28^ showed either no or very little depression following the first stimulus, while STX1A^ΔN2-9^ exhibited less depression than STX1A^WT^ after the first 10 stimuli (Fig 6A).

**Figure 6:**
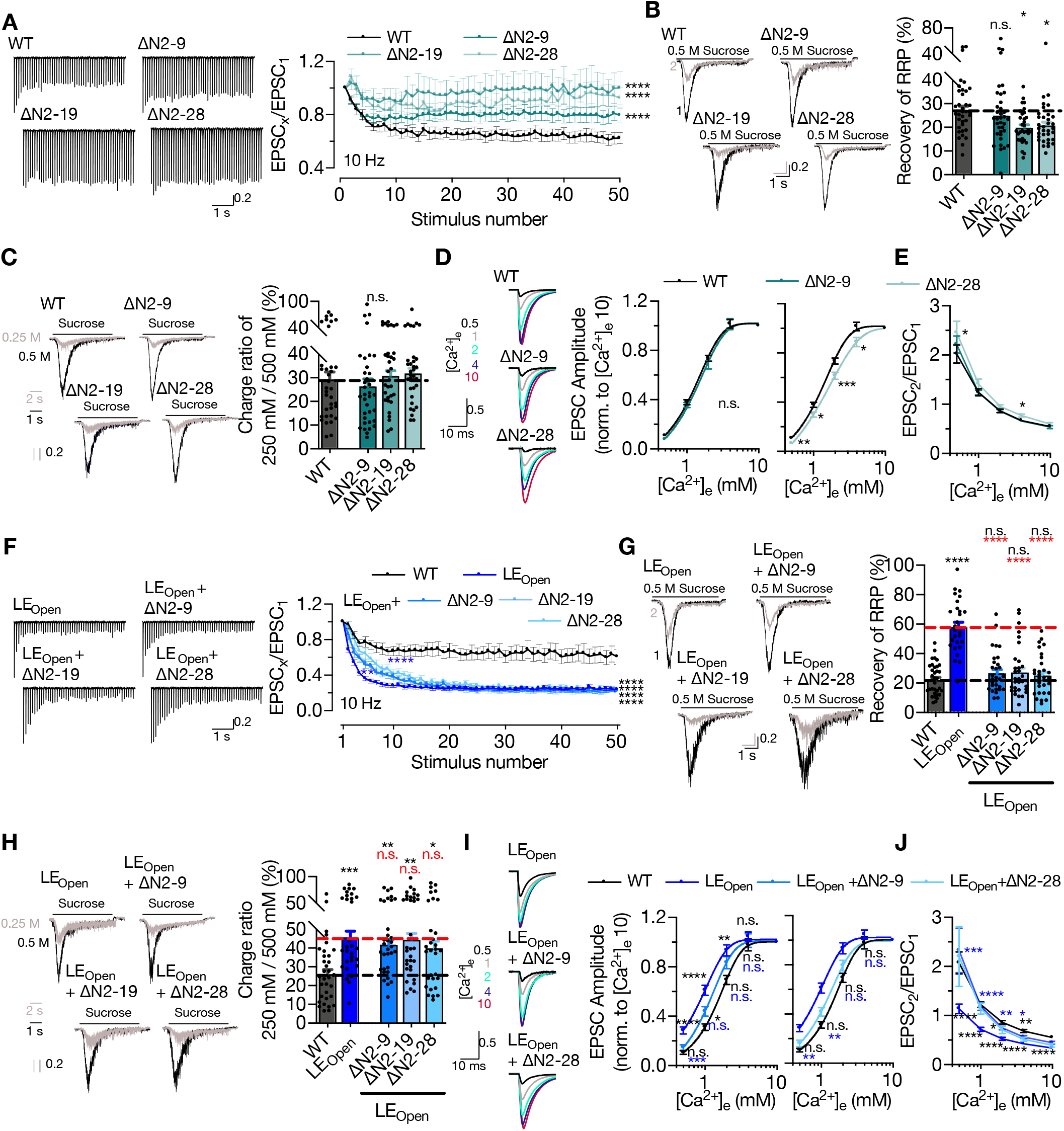
STX1’s N-peptide has a modulatory function in short-term plasticity and Ca^2+^-sensitivity of synaptic transmission. A. Example traces (left) and quantification (right) of STP measured by 50 stimulations at 10 Hz from STX1A^WT^, STX1A^ΔN2-9^, STX1A^ΔN2-19^, or STX1A^ΔN2-28^ neurons. The responses were normalized to the EPSC_1_. B. Example traces (left) and quantification (right) of the recovery of RRP determined as the fraction of RRP measured at a second pulse of 500 mM sucrose solution after 2s of initial depletion from STX1A^WT^, STX1A^ΔN2-9^, STX1A^ΔN2-19^, or STX1A^ΔN2-28^ neurons. C. Example traces (left) and quantification (right) of the ratio of the charge transfer triggered by 250 mM sucrose over that of 500 mM sucrose as a read-out of fusogenicity of the SVs. D. Example traces (left) and quantification (right) of Ca^2+^-sensitivity as measured by the ratio of EPSC amplitudes at [Ca^2+^]_e_ of 0.5, 1, 2, 4, and 10 mM recorded from STX1A^WT^, STX1A^ΔN2-9^, or STX1A^ΔN2-28^ neurons. The responses were normalized to the response at [Ca^2+^]_e_ of 10 mM. E. PPR of EPSC amplitudes at [Ca^2+^]_e_ of 0.5, 1, 2, 4, and 10 mM recorded at 40 Hz. F. Example traces (left) and quantification (right) of STP measured from STX1A^WT^, STX1A^LEOpen^, STX1A^LEOpen + ΔN2-9^, STX1A^LEOpen + ΔN2-19^, or STX1A^LEOpen + ΔN2-28^ neurons. G. Example traces (left) and quantification (right) of the recovery of RRP 2 s after initial depletion. H. Example traces (left) and quantification (right) of the ratio of the charge transfer triggered by 250 mM sucrose over that of 500 mM sucrose. I. Example traces (left) and quantification (right) of Ca2+-sensitivity recorded from STX1A^WT^, STX1A^LEOpen^, STX1A^LEOpen^ ^+ ΔN2-9^, or STX1A^LEOpen^ ^+ ΔN2-28^ neurons. The responses were normalized to the response at [Ca^2+^]_e_ of 10 mM. J. PPR of EPSC amplitudes at [Ca^2+^]_e_ of 0.5, 1, 2, 4, and 10 mM recorded at 40 Hz. Data information: The artifacts are blanked in example traces in (A, D, F, & I). In (A, D-F, I & J), data points represent the mean ± SEM. In (B, C, G &, H), data points represent single observations, the bars represent the mean ± SEM. Black and red annotations on the graphs show the significance comparisons to STX1A^WT^ or STX1A^LEOpen^ neurons, respectively (either nonparametric Kruskal-Wallis followed by Dunn’s *post hoc* test or one-way ANOVA followed by Holm Sidak’s *post-hoc* test was applied based on the normality of the data. Two-way ANOVA was applied for data in (A) and (F), *p ≤ 0.05, **p ≤ 0.01, ***p ≤ 0.001, ****p ≤ 0.0001). The numerical values are summarized in source data.

Whereas Pvr shapes the STP curve starting from the initial phase, the late phase of STP is affected not only by Pvr, but also by the rate of SVs newly arriving at the AZ. Because all the STX1A^ΔN^ neurons showed an altered behavior in the late phase of STP, we hypothesized that these neurons might keep up better with the high-frequency stimulus than STX1A^WT^ does because of an increase in newly arrived SVs, or replenishment of the RRP. To study the efficacy of replenishment of the primed vesicles, we stimulated the neurons with a double pulse of 500 mM sucrose solution with a time interval of 2 s (Fig 6B), as the replenishment of the whole pool of the primed vesicles after sucrose depletion takes at least 10 s (Stevens & Tsujimoto, 1995). STX1A^ΔN2-9^ showed no effect on the fraction of the RRP recovered after full depletion when compared to that of STX1A^WT^ (Fig 6B). On the other hand, STX1A^ΔN2-19^ and STX1A^ΔN2-19^ significantly decreased the vesicle replenishment rate by ~10%, which is contrary to our initial expectation (Fig 6B). This suggests that an increased replenishment rate does not account for the decreased depression in the STP curves of STX1A^ΔN^ mutants.

Pvr and the degree of STP depend on both vesicle fusogenicity and Ca^2+^-sensitivity of the vesicular release, thus we scrutinized the effects of N-peptide deletions on these variables. Calculating the RRP fraction released in response to a subsaturating, 250 mM sucrose solution application revealed no difference between STX1A^WT^ and STX1A^ΔN^ neurons, suggesting no decrease in SV fusogenicity with these mutants (Fig 6C). Interestingly, when we generated Ca^2+^-dose response curves by evoking AP driven EPSCs in the presence of 0.5, 1, 2, 4, or 10 mM Ca^2+^-containing extracellular solutions, we observed a lowered apparent Ca^2+^-sensitivity in the STX1A^ΔN2-19^ (Fig EV4) and STX1A^ΔN2-28^ neurons (Fig 6D) when compared to STX1A^WT^ neurons. On the other hand, STX1A^ΔN2-9^ neurons showed a normal pattern of increase in EPSCs in relation to increasing extracellular Ca^2+^-concentration (Fig 6D). We also measured the PPR, which is inversely related to Pvr, at different extracellular Ca^2+^-concentrations and determined that STX1A^ΔN2-28^ had a significantly higher PPR at 0.5 mM [Ca^2+^]_e_ compared to that of STX1A^WT^ (Fig 6E). This was also evident in STP behavior elicited by 5 AP stimulation at 40 Hz, as STX1A^ΔN2-28^ neurons showed a greater facilitation at 0.5 mM [Ca^2+^]_e_ compared to that of STX1A^WT^ (Fig EV4). Whereas increasing [Ca^2+^]_e_ to 2mM was not sufficient to drive the STP behavior of STX1A^ΔN2-28^ neurons towards STX1A^WT^-like pattern, at the highest [Ca^2+^]_e_ tested all the groups STX1A^WT^, STX1A^ΔN2-9^, and STX1A^ΔN2-28^ showed a similar level of depression upon 40 Hz stimulation (Fig EV4).

It is well documented that the presumed opening of STX1 and thus the increase in Pvr by LE_Open_-mutation enhances short-term depression. Deletion of N-peptide at any length in STX1A^LEOpen^ did not change the degree of the depression in the late phase of the high-frequency stimulus, however all the deletions decreased the slope of the depression in the initial phase compared to that of STX1A^LEOpen^ alone (Fig 6F). The recovery of the RRP after sucrose depletion was enhanced by the open conformation of STX1A but reverted back to the WT-like levels by the expression of STX1A^LEOpen+ΔN^ mutants (Fig 6G). Strikingly, the increase in fusogenicity was not influenced by the N-peptide deletions (Fig 6H). On the other hand, N-peptide deletions imposed a right-shift in the Ca^2+^-dose response curves on the STX1A^LEOpen^, which markedly increased the Ca^2+^-sensitivity, making them approach again WT-like levels (Fig 6I & Fig EV4). PPR was also comparable between STX1A^WT^ and STX1A^LEOpen+ΔN^ mutants, whereas STX1A^LEOpen^ always showed a greatly reduced PPR in all extracellular Ca^2+^-concentrations tested (Fig 6J). Contrary to STX1A^WT^ neurons, STX1A^LEOpen^ neurons showed no facilitation at 0.5 mM [Ca^2+^]_e_, and a greater depression at 2 mM [Ca^2+^]_e_ as well as at 10 mM [Ca^2+^]_e_. Whereas STX1A^LEOpen+ΔN2-28^ showed a similar pattern of facilitation at 0.5 mM [Ca^2+^]_e_ to STX1A^WT^, at higher [Ca^2+^]_e_ their short-term-depression approached the level of STX1A^LEOpen^ (Fig S6).

Decreased Ca^2+^-sensitivity can arise from either reduced Ca^2+^-influx as a result of alterations in Ca^2+^-channel localization or gating, or from a disturbance in Ca^2+^-secretion coupling. To address that issue, we expressed the Ca^2+^-reporter GCamp6f coupled to Synaptophysin (SynGCamp6f) in STX1-null neurons with or without STX1A rescue constructs and measured the immunofluorescence at the synapses at baseline or upon 1, 2, 5, 10, or 20 AP stimulation at 10 Hz (Fig 7A-7E). Surprisingly, STX1-null neurons showed a decreased Ca^2+^-influx compared to neurons rescued with STX1A^WT^ (Fig 7C). However, STX1A^ΔN2-9^ or STX1A^ΔN2-28^ did not influence the SynGCamp6f signal at any AP number elicited (Fig 7D), whereas Ca^2+^-influx was reduced in synapses in STX1A^LEOpen^ and STX1A^LEOPen+ΔN2-28^ neurons (Fig 7E).

**Figure 7:**
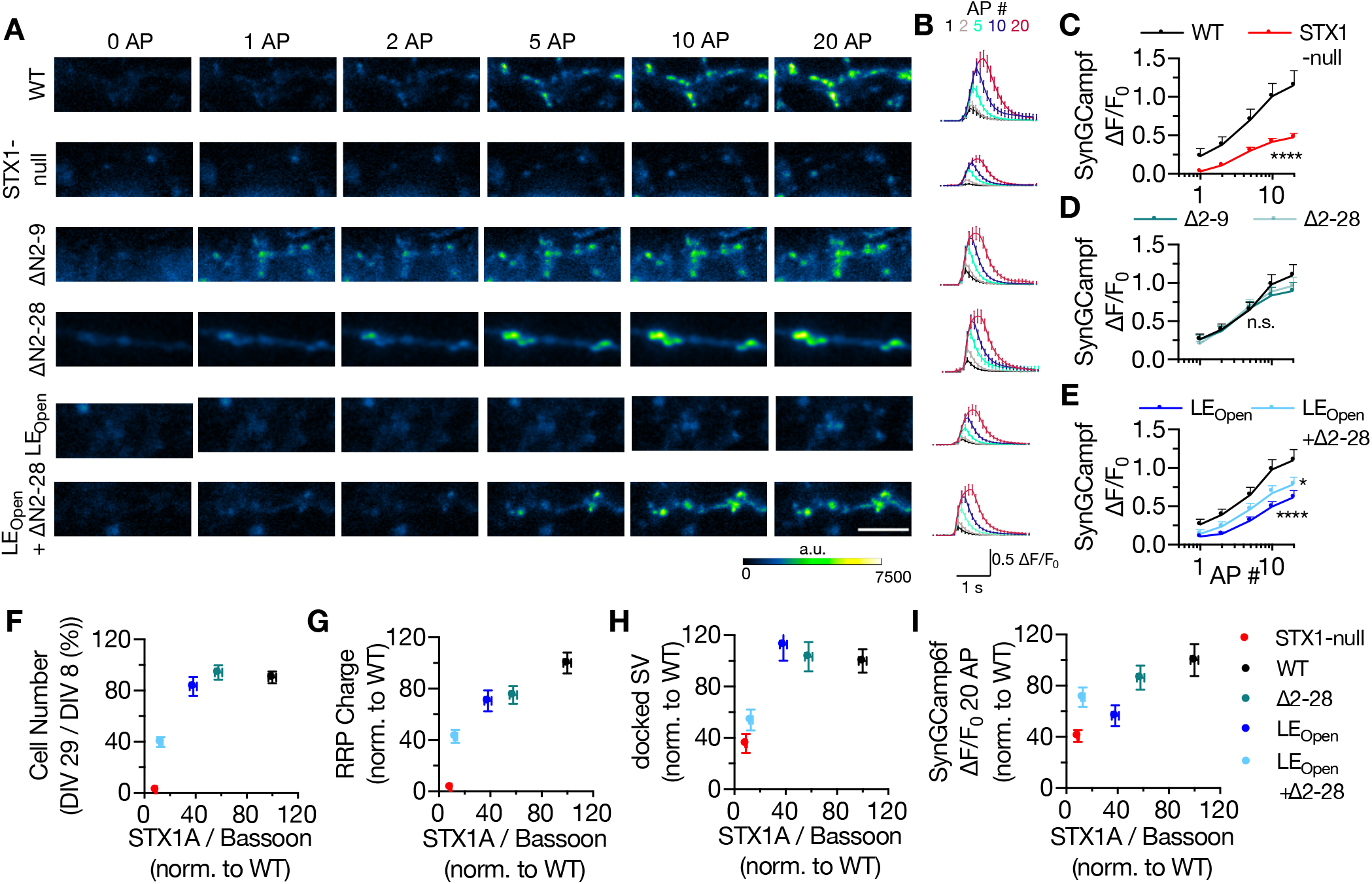
Ca^2+^-influx is reduced in STX1-null and in STX1^LEOpen^ neurons. A-B. Example images and average of SynGCaMP6f fluorescence as (ΔF/F_0_) in STX1-null neurons either not rescued or rescued with STX1A^WT^, STX1A^ΔN2-9^, STX1A^ΔN2-19^, STX1A^LEOpen^, or STX1A^LEOpen^ ^+ ΔN2-28^. The images were recorded at baseline, and at 1, 2, 5, 10, and 20 APs. Scale bar: 10 μm C-D-E. Maximum fluorescence changes (ΔF/F_0_) in STX1-null, STX1A^ΔN2-9^, STX1A^ΔN2-28^, STX1A^LEOpen^, or STX1A^LEOpen^ ^+ ΔN2-28^. In comparison to that in STX1A^WT^ neurons recorded at 1, 2, 5, 10, and 20 APs. F-G-H-I. Plot of neuronal viability and RRP charge, number of docked SVs and maximum SynGCaMP6f ΔF/F_0_ at 20 AP from STX1-null, STX1A^WT^, TX1A^ΔN2-28^, STX1A^LEOpen^, and STX1A^LEOpen^ ^+ ΔN2-28^ normalized to STX1A^WT^ neurons as a function of STX1A expression. All the values were normalized to the one obtained from STX1A^WT^ neurons in each individual culture. Data information: Data points in all graphs represent the mean ± SEM. Black annotations on the graphs show the significance comparisons to STX1A^WT^ (either unpaired t-test or Mann-Whitney test was applied in (C) based on the normality of the data. In (D) and (E) nonparametric Kruskal-Wallis test followed by Dunn’s *post hoc* test was applied, *p ≤ 0.05, ****p ≤ 0.0001). The numerical values are summarized in source data.

The reduction in Ca^2+^-influx at the presynaptic terminals in STX1-null, STX1A^LEOpen^ and STX1A^LEOPen+ΔN2-28^ neurons compared to that of STX1A^WT^ neurons is indicative of involvement of STX1 in the vesicular release processes upstream of vesicle docking (Fig 7A-7E). As these STX1A mutants also showed severely decreased expression levels (Fig 4), we hypothesized that the structural properties as well as the neuronal viability might be affected by the expression level of STX1A. As a potential correlation of STX1A’s function to its expression level, we plotted the neuronal viability, the size of the RRP, the number of docked SV, or the level of Ca^2+^-influx at 20 AP normalized to the values obtained from STX1A^WT^ neurons as a function of STX1A expression and observed all the parameters showed a decreased degree of rescue with decreased levels of STX1A (Fig 7F-7I).

## Discussion

The tight interaction between STX1 and Munc18-1 is not dictated through a single contact point but rather spans a large area both on STX1 and Munc18-1 (Misura *et al.*, 2000), which are mostly driven by STX1’s N-peptide and closed conformation. Using our STX1-null mouse model system, we can draw several conclusions from mutant STX1 rescue experiments: (1) STX1’s H_abc_-domain is essential for the stability of STX1 and Munc18-1, and thus for neurotransmitter release and overall STX1 function; (2) STX1’s N-peptide is dispensable for neurotransmitter release, but has a minor modulatory function for STX1’s stability and for Ca^2+^-sensitivity of vesicular release; and (3) neurotransmitter release can proceed even when both interaction modes are presumably intervened by N-peptide deletions in conjunction with LE_Open_-mutation in STX1A and even when both proteins are expressed at severely low levels.

### STX1’s H_abc_-domain is essential for the overall function of STX1

The three helical H_abc_-domain constitutes a major portion of STX1. As the main driving force for vesicle fusion is the zippering of the SNARE domains of STX1, SNAP25, and Syb2 (Baker & Hughson, 2016; Rizo & Sudhof, 2012; Rizo & Xu, 2015), STX1, which lacks only the H_abc_-domain sufficiently mediates liposome fusion in reconstitution experiments (Rathore *et al.*, 2010; Shen *et al.*, 2010). Even in the synaptic environment, a crucial function of the H_abc_-domain has been suggested only for spontaneous neurotransmitter release (Meijer *et al.*, 2012; Zhou *et al.*, 2013). However, the picture in intact synapses is more complex, because neurotransmitter release proceeds as a result of multiple steps dependent on protein folding and trafficking, intermolecular interactions, relative conformations, and the proper localization of multiple synaptic proteins.

What is clear from our study and the previous studies is the importance of H_abc_-domain in proper folding of STX1 and its co-recruitment to the AZ with Munc18-1. Severely decreased expression levels of STX1, Munc18-1, or both occur when the interaction between the H_abc_-domain of STX1 and Munc18-1 is interrupted (Gulyas-Kovacs *et al.*, 2007; Meijer *et al.*, 2012; Vardar *et al.*, 2020; Zhou *et al.*, 2013). Even improper folding of the H_abc_-domain by an insertion/deletion (InDel) mutation, identified in relation to epilepsy, leads to a high degree of STX1 instability (Vardar *et al.*, 2020). Consistently, both the STX1B InDel mutant (Vardar *et al.*, 2020) and STX1A^ΔHabc^ mutant (Fig. 2) were incapable of sustaining neuronal viability. Due to the lack of significant expression of STX1A^ΔHabc^ (Fig 1 & EV1), we cannot draw a certain conclusion on whether or not H_abc_-domain of STX1 is directly involved in neurotransmitter release. Yet, we argue that the major role of H_abc_-domain of STX1 is to drive it into its correct folding and to recruit it together with Munc18-1 to the AZ.

### STX1’s N-peptide role in neurotransmitter release is only detectable in STX1’s open configuration

On the contrary to the general view, we show here that the STX1’s N-peptide is not indispensable for neurotransmitter release, but rather only modulates STX1’s expression and the Ca^2+^-sensitivity of SVs. Above all, the dispensability of STX1’s N-peptide in vesicle fusion and particularly for proper recruitment of Munc18-1 to the AZ is consistent with the estimated contribution of N-peptide to the overall affinity of STX1 to Munc18-1, which is only minor (Burkhardt *et al.*, 2008; Christie *et al.*, 2012; Colbert *et al.*, 2013).

Remarkably, the putative loss of STX1–Munc18-1 as a binary complex by interference with both canonical interaction modes has little or no effect on synaptic transmission in general. Is it possible that STX1 and Munc18-1 only ensure the stability of one another and that a permanent interaction between these proteins is not required for vesicle fusion? Besides showing a largely unaltered affinity for Munc18-1, STX1A^ΔN^ mutants and STX1A^LEOpen^ have also been proposed to maintain the closed conformation when bound to Munc18-1 (Colbert *et al.*, 2013; Dawidowski & Cafiso, 2013), which also includes contact points to STX1’s SNARE motif (Burkhardt *et al.*, 2008; Liang *et al.*, 2013; Misura *et al.*, 2000). Given that and the flexibility of STX1-Munc18-1 interaction, which induces large conformational changes on these proteins not only when STX1 is isolated but also when it enters the SNARE complex (Jakhanwal *et al*, 2017), it is conceivable that even for the STX1A^LEOpen+ΔN^ mutants a level of interaction between STX1–Munc18-1 must be retained.

Nevertheless, our analysis shows that even though both N-peptide deletion and LE_Open_-mutation produce the same degree of reduction in the binding affinity of STX1 to Munc18-1 (Burkhardt *et al.*, 2008; Christie *et al.*, 2012; Colbert *et al.*, 2013), it is the closed conformation that commands the Munc18-1 recruitment and/or stability at the synapse (Fig.1). This led us to interpret Munc18-1’s binding to STX1 and its ultimate effect on SNARE complex formation as a two-step process, which is a convolution of the affinity and the efficacy of this interaction. Consistently, it has been thought that LE_Open_-mutation exposes the SNARE domain of STX1 (Dulubova *et al.*, 1999), whereas absence of N-peptide tightens its Munc18-1 driven closed conformation (Christie *et al.*, 2012; Colbert *et al.*, 2013; Khvotchev *et al.*, 2007) potentially resulting in opposing effects in SNARE complex formation. Indeed, our observation that N-peptide deletion reverses the STX1A^LEOpen^-dependent facilitation of neurotransmitter release parameters, which is generally attributed to its promotion of SNARE complex formation (Acuna *et al*; Dulubova *et al.*, 1999; Gerber *et al.*, 2008), hints at a reduction in the number of SNARE complexes formed by STX1A^LEOpen^. However, N-peptide likely plays only a minor role in determining the equilibrium of open-closed conformations of STX1^WT^ in a membranous environment when STX1’s TMR is present (Dawidowski & Cafiso, 2013), and thus the modulation of neurotransmitter release by the N-peptide cannot be observed in STX1^WT^ but can be only unmasked by the introduction of LE_Open_-mutation.

Importantly, Munc18-1 does not bind only to STX1, but it is also thought to bind to the SNARE complex formed by STX1, SNAP-25, and Syb-2 (Burkhardt *et al.*, 2008; Dulubova *et al*, 2007; Shen *et al.*, 2007; Zilly *et al*, 2006) to provide a template as a scaffold together with Munc13 (Ma *et al*, 2013; Ma *et al*, 2015). Given that the SNARE complex formation is a dynamic process, which involves not only the assistance but also the protection by Munc18-1 against NSF-dependent dissociation (He *et al*, 2017), such a two-step process is also applicable for the back- and forward shift in the number of SNARE complexes. Thus, it is plausible that the stability of the SNARE complex is ensured by Munc18-1’s efficient binding to STX1, which may account for the additive effects of N-peptide deletion and LE_Open_-conformation on the size of RRP (Fig 4 & Fig 7). However, this regulation process must be upstream of the vesicle priming, as the fusogenicity of the primed vesicles was predominantly dictated by the LE_Open_-mutation.

How the synaptic vesicles become more fusogenic in the presence of STX1A^LEOpen^ is not known, though one simple explanation is its propensity to produce reactive SNARE complexes with a higher number and efficacy (Acuna *et al.*, 2014; Dulubova *et al.*, 1999). This hypothesis is appealing as it can also account for the faster recovery of the SVs to the RRP by LE_Open_-mutation (Fig 6). Surprisingly, however, addition of N-peptide deletions not only in STX1A^LEOpen^ but also in STX1A^WT^ exclusively slowed down the RRP replenishment without an effect on the SV fusogenicity (Fig 6), uncoupling the regulation of these two processes. We cannot explain this phenomenon based on our data but would like to draw attention to that there are still unsolved questions regarding the regulation of SV fusogenicity. In fact, it is thought that at the state of primed and even docked vesicles the SNAREs are zippered up to hydrophobic layer +4 (Sorensen *et al*, 2006; Vardar *et al.*, 2016; Walter *et al*, 2010) and thus include already ‘opened’ STX1. Therefore, the increase in SV fusogenicity by STX1A^LEOpen^- and STX1A^LEOpen+ΔN^-mutants might involve yet an unknown mechanism, which does not employ STX1’s N-peptide.

On the other hand, it is remarkable that the deletion of the N-peptide of 19 or 28 aa reduced the Ca^2+^-sensitivity of the vesicular release both in default and open STX1A. Ca^2+^-sensitivity of the vesicular release is estimated by the assessment of Ca^2+^-dose response, which is a convoluted measurement of SV fusogenicity, Ca^2+^-influx, and Ca^2+^-channel–SV distance coupling. Accordingly, the increased Ca^2+^-sensitivity of vesicular release by LE_Open_-mutation stems partly from the increased fusogenicity of SVs (Fig 6). However, the rightward shift in Ca^2+^-dose response curve caused by N-peptide deletions was not accompanied by an altered fusogenicity of SVs neither in default or open configuration of STX1A. Thus, it is conceivable that the N-peptide deletions might have led to disrupted Ca^2+^-channel–SV coupling. This is also consistent with that the STX1ALEOpen neurons have an enhanced Ca^2+^-dependent release (Fig. 6), yet they manifest a reduction in global Ca^2+^-influx at the presynaptic terminals (Fig. 7). In fact, a direct interaction between STX1 and Ca^2+^-channels (Bachnoff *et al*, 2013; Cohen *et al*, 2007; Sheng *et al*, 1994; Wiser *et al*, 1996), has been proposed to contribute to the overall Ca^2+^-sensitivity of the vesicular release machinery, where STX1 deemed an inhibitory role in baseline activity of Ca^2+^-channels (Trus *et al*, 2001). Additionally, this suggests a regulatory function of STX1 independent of Munc18-1, which is in line with unaltered STP behavior upon expression of Munc18-1 mutant, which cannot bind to the STX1’s N-peptide (Meijer *et al.*, 2012). However, our finding that Ca^2+^-influx was not affected by N-peptide deletions challenges this hypothesis.

Alternatively, the CK2 binding motif 14-SDDDDD-19 on STX1 could act as a secondary Ca^2+^-sensor for neurotransmitter release, yet this hypothesis also falls short as the shortest deletion of N-peptide aa 2-9 also reduces the Ca^2+^-sensitivity of the vesicles when introduced in the STX1A^LEOpen^ without a significant effect on vesicle fusogenicity. Therefore, we argue that the most likely explanation of the reduced Ca^2+^-sensitivity is again the possibly reduced number of SNARE complexes, consistent with the hypothesis that the increased Ca^2+^-sensitivity by STX1^LEOpen^ is due to the increased number of SNARE complexes (Acuna *et al.*, 2014). An impairment in SNARE complex formation additionally could explain the slower rate of the recovery of the RRP in neurons that express STX1A^ΔN2-19^ or STX1A^ΔN2-28^.

## Material and Methods

### Animal maintenance and generation of mouse lines

All procedures for animal maintenance and experiments were in accordance with the regulations of and approved by the animal welfare committee of Charité Medical University and the Berlin state government Agency for Health and Social Services under license number T0220/09. The generation of STX1-null mouse line was described previously (Arancillo *et al.*, 2013; Vardar *et al.*, 2016).

### Neuronal cultures and lentiviral constructs

Hippocampal neurons were obtained from mice of either sex at postnatal day (P) 0–2 and seeded on the already prepared continental or micro-island astrocyte cultures as described previously (Vardar *et al.*, 2016; Xue *et al*, 2007). The neuronal cultures were then incubated for 13-20 DIV in Neurobasal™-A supplemented with B-27 (Invitrogen), 50 IU/ml penicillin and 50 μg/ml streptomycin at 37°C before experimental procedures. Neuronal cultures for EM and Ca^2+^-influx and those for neuronal viability, immunofluorescence labeling, and electrophysiology experiments were transduced with lentiviral particles at DIV 2-3 and DIV 1, respectively. Lentiviral particles were provided by the Viral Core Facility (VCF) of the Charité-Universitätsmedizin, Berlin, and were prepared as previously described (Vardar *et al.*, 2016).. The cDNA mouse STX1A (NM_016801) was cloned in frame after an NLS-GFP-P2A sequence. The improved *Cre* recombinase (iCre) cDNA was fused to NLS-RFP-P2A. SynGCamp6f was generated analogous to synGCamp2 (Herman *et al*, 2014), by fusing GCamp6f (Chen *et al*, 2013) to the C terminus of synaptophysin and cloned also in frame after an NLS-GFP-P2A sequence.

### Neuronal viability

The *in vitro* viability of the neurons was defined as the percentage of the number of neurons alive at DIV15, 22, 29, 36, and 43 compared to the number of neurons at DIV 8. Phase contrast bright field images and fluorescent images with excitation wavelengths of 488 and 555 nm were acquired with a DMI 400 Leica microscope, DFC 345 FX camera, HCX PL FLUOTAR 10 objectives, and LASAF software (all from Leica). Fifteen randomly selected fields of 1.23 mm^2^ per well and two wells per group in each culture were imaged at different time points and the neurons were counted offline with the 3D Objects Counter function in Fiji software as described previously (Vardar *et al.*, 2016). MAP2 immunofluorescence labelling as shown in the figures is used only for representative purposes.

### Immunocytochemistry

The neuronal cultures were fixed and immunostained as described previously (Vardar *et al.*, 2016) and in Appendix Supplementary Methods. The primary and secondary antibodies are listed in the Appendix Supplementary Methods. The images were acquired with an Olympus IX81 epifluorescent-microscope with MicroMax 1300YHS camera using MetaMorph software (Molecular Devices) and analyzed offline with ImageJ as previously described (Vardar *et al.*, 2016).

### Electrophysiology

Whole cell patch-clamp recordings were performed on glutamatergic autaptic hippocampal neurons at DIV 14–20 at RT with a Multiclamp 700B amplifier and an Axon Digidata 1550B digitizer controlled by Clampex 10.0 software (both from Molecular Devices), as described previously (Vardar *et al.*, 2020) and in Appendix Supplementary Methods. The recordings were analyzed offline using Axograph X Version 1.7.5 (Axograph Scientific).

Additionally, for Ca^2+^-sensitivity assays, 6 EPSCs were evoked at 0.2 Hz in extracellular solution containing 1 mM MgCl_2_ and either 0.5, 1, 2, 4, or 10 mM CaCl_2_. In between the test extracellular solution applications, standard extra cellular solution was applied to control for rundown and cell-to-cell variability and the responses in text solutions were normalized to the preceding responses in standard solution. The then normalized values to those in 10 mM CaCl_2_ were fitted into a standard Hill equation.

### SynGcamp6f-imaging

Imaging experiments were performed at DIV 13-16 on autapses in response to a single stimulus and stimuli trains of 10 Hz. Images were acquired using a 490-nm LED system (pE2; CoolLED) at a 5 Hz sampling rate with 25 ms of exposure time. The acquired images were analyzed offline using ImageJ (National Institute of Health), Axograph X (Axograph), and Prism 8 (Graph-Pad; San Diego, CA).

### EM

The high-density cultured hippocampal neurons high-pressure fixed using an HPM 100 Leica or ICE Leica high-pressure freezer at DIV 14-16 as described previously (Vardar *et al.*, 2016) and in Appendix Supplementary Methods. Synapses were analyzed blindly using an analysis program developed for ImageJ and MATLAB as described previously (Watanabe *et al*, 2013).

### Statistical analysis

Data in bar graphs present single observations (points) and means ± standard error of the mean (SEM; bars). Data in x-y plots present means ± SEM. All data were tested for normality with Kolmogorov-Smirnov test. Data from two groups with normal or non-parametric distribution were subjected to Student’s two-tailed t-test or Mann-Whitney non-parametric test, respectively. Data from more than two groups were subjected to Kruskal-Wallis followed by Dunn’s *post hoc* test when at least one group showed a non-parametric distribution. For data in which all the groups showed a parametric distribution, one-way ANOVA test followed by Tukey’s *post hoc* test was applied. For STP measurements, two-way ANOVA test was used. All the tests were run with GraphPad Prism 8.3 and all the statistical data are summarized in corresponding source data tables.

## Acknowledgements

We thank the Charité Viral Core facility, Katja Pötschke, and Bettina Brokowski for virus production, Berit Söhl-Kielczynski and Heike Lerch for technical assistance, Melissa Herman for her contribution to the manuscript and to all the Rosenmund Lab members for the discussions. This work was supported by the German Research Council (Collaborative Research Grant SFB958, TRR186, and Reinhart Koselleck Project).

## Author contributions

GV and CR designed the experiments and interpreted the data. GV performed immunostainings and imaging, neuronal survivability assays, electrophysiology and Ca^2+^-imaging experiments and analyzed the obtained data and the electron microscopy data. ASL, VZ, and MWB performed electrophysiology experiments and analyzed the data. MWB obtained Ca^2+^-imaging data. MB obtained electron microscopy data. SZ performed neuronal survivability assays and analyzed the data. TT designed the lentiviral constructs. GV designed the figures. GV and CR wrote the manuscript.

**Figure EV1: STX1A^ΔHabc^ expression cannot be detected**

A. Example images (left) and quantification of immunofluorescence labeling for Bassoon and STX1A shown as red and green, respectively, in the corresponding composite pseudocolored images obtained from high-density cultures of STX1-null hippocampal neurons either not rescued or rescued with STX1A^WT^, or STX1A^WT-FLAG^. The data were normalized to the STX1A fluorescence/Bassoon fluorescence value obtained from STX1A^WT^ neurons. Scale bar: 10 μm

A. Example images (left) and quantification of immunofluorescence labeling for Bassoon and FLAG shown as red and green, respectively, in the corresponding composite pseudocolored images obtained from high-density cultures of STX1-null hippocampal neurons rescued with either STX1A^WT^, STX1A^WT-FLAG^, STX1A^ΔN2-9-FLAG^, STX1A^LEOpen-FLAG^, or STX1A^ΔHabc-FLAG^. The data were normalized to the STX1A fluorescence/Bassoon fluorescence value obtained from STX1A^WT-FLAG^ neurons. Scale bar: 10 μm

Data information: Data points in graphs represent single observations, and the bars represent the mean ± SEM. In (A), black and red annotations on the graphs show the significance comparisons to STX1-null and STX1A^WT^, respectively. In (B), black and red annotations on the graphs show the significance comparisons to STX1^WT^ and STX1A^WT-FLAG^, respectively (nonparametric Kruskal-Wallis test followed by Dunn’s *post hoc* test, *p ≤ 0.05, **p ≤ 0.01, ***p ≤ 0.001, ****p ≤ 0.0001). The numerical values are summarized in Appendix Table 1.

**Figure EV2: Reducing the expression levels of STX1A^WT^ or STX1A^LEOpen^ does not alter their synaptic release properties**

A. Example images of immunofluorescence labeling for VGlut1, STX1A, Munc18-1, and NLS-GFP shown as red, green, blue, and gray, respectively, in autaptic STX1-null hippocampal neurons rescued with 1X viral volume of STX1A^LEOpen^ ^+ ΔN2-28^ as reference or with different viral volumes of either STX1A^WT^, or STX1A^LEOpen^ as. Scale bar: 10 μm

B. Higher magnification images of the areas highlighted in the images in **A**. VGlut1, STX1A, and Munc18-1 are shown as red, green, and blue, respectively in the corresponding composite image. Scale bar: 10 μm

C. Quantification of the fluorescence intensity of NLS-GFP from the neurons as in **A** as normalized to that of STX1A^WT^ 1X rescue.

D. Quantification of the immunofluorescence intensity of STX1A as normalized to the immunofluorescence intensity of Bassoon in the same ROIs as shown in **B**. The values were then normalized to the values obtained from STX1A^WT^ 1X neurons.

E. Quantification of the immunofluorescence intensity of Munc18-1 as normalized to the immunofluorescence intensity of Bassoon in the same ROIs as shown in **B**. The values were then normalized to the values obtained from STX1-null neurons expressing STX1A^WT^ 1X neurons.

F. Example traces (left) and quantification of the amplitude (right) of EPSCs obtained from hippocampal autaptic STX1-null neurons rescued with 1X viral volume of STX1A^LEOpen^ ^+ ΔN2-28^ as reference or with different viral volumes of either STX1A^WT^, or STX1A^LEOpen^ as. The artifacts are blanked in the example traces.

G. Example traces (left) and quantification of the charge transfer (right) of sucrose-elicited RRPs obtained from the same neurons as in **F**.

H. Quantification of Pvr determined as the percentage of the RRP released upon one action potential.

I. Example traces (left) and quantification of the frequency (right) of mEPSCs recorded at −70 mV. The example traces were filtered at 1 kHz.

Data information: In (C–I), data points represent single observations, the bars represent the mean ± SEM. Red annotations (stars and n.s.) on the graphs show the significance comparisons to STX1A^LEOpen^ ^+ ΔN2-28^ 1X rescue. Black annotations (stars and n.s.) on the bars show the significance comparisons for rescues using different viral volume either for STX1A^WT^ or STX1A^LEOpen^. (nonparametric Kruskal-Wallis test followed by Dunn’s *post hoc* test, *p ≤ 0.05, **p ≤ 0.01, ***p ≤ 0.001, ****p ≤ 0.0001). The numerical values are summarized in Appendix Table 2.

**Figure EV3: Interruption of both Munc18-1 binding modes of STX1 ultimately leads to neuronal death**

A. Example images of high-density cultures of STX1-null hippocampal neurons at DIV 8, 15, 22, 29, 36, and 43 represented with immunofluorescent labeling of MAP2. Red and green nuclei serve as a marker for NLS-RFP-P2A-*Cre* recombinase expression and for NLS-GFP-P2A-STX1A (either WT or mutants), respectively. 50 μm.

B. Quantification of neuronal density at DIV 8 in STX1-null hippocampal neurons either not rescued or rescued with STX1A^WT^, STX1A^LEOpen^, STX1A^LEOpen + ΔN2-9^, STX1A^LEOpen + ΔN2-19^, or STX1A^LEOpen + ΔN2-28^. Scale bar: 10 μm

C. Quantification of the percentage of the surviving neurons at DIV 8, 15, 22, 29, 36, and 43 as normalized to the neuronal density at DIV 8.

Data information: In (B), data points represent single observations, the bars represent the mean ± SEM. In (C), data points represent the mean ± SEM. Red and black annotations (stars and n.s.) on the graphs show the significance comparisons to STX1-null and to STX1A^WT^ rescue, respectively (nonparametric Kruskal-Wallis test followed by Dunn’s *post hoc* test, *p ≤ 0.05, **p ≤ 0.01, ***p ≤ 0.001, ****p ≤ 0.0001). The number of observations and cultures and statistical data are summarized in source data. The numerical values are summarized in Appendix Table 3.

**Figure EV4: Deletion of N-peptide increases the PPR both in closed and open conformation of STX1A in low extracellular Ca^2+^-concentration**

A. Example traces (left) and quantification (right) of Ca^2+^-sensitivity recorded from neurons expressing STX1A^ΔN2-19^, compared to that of neurons expressing STX1A^WT^. The responses were normalized to the response at [Ca^2+^]_e_ of 10 mM. The artifacts are blanked in the example traces.

B. Example traces (left) and quantifications (right) of STP measured at [Ca^2+^]_e_ of either 0.5 mM, 2 mM, or 10 mM by 5 stimulations at 40 Hz from neurons expressing either STX1A^WT^, STX1A^ΔN2-9^, or STX1A^ΔN2-28^. The artifacts are blanked in the example traces. At every [Ca^2+^]_e_, the responses were normalized to the EPSC_1_.

C. Example traces (left) and quantification (right) of Ca^2+^-sensitivity recorded from neurons expressing STX1A^LEOpen^ ^+ ΔN2-19^, compared to that of neurons expressing STX1A^WT^ or STX1A^LEOpen^. The responses were normalized to the response at [Ca^2+^]_e_ of 10 mM. The artifacts are blanked in the example traces.

D. Example traces (left) and quantifications (right) of STP measured at [Ca^2+^]_e_ of either 0.5 mM, 2mM, or 10mM by 5 stimulations at 40 Hz from neurons expressing either STX1A^WT^, STX1A^LEOpen^ ^+ ΔN2-9^, or STX1A^LEOpen^ ^+ ΔN2-28^. The artifacts are blanked in the example traces. At every [Ca^2+^]_e_, the responses were normalized to the EPSC_1_.

Data information: In (A-D), data points represent the mean ± SEM. Black and blue annotations (stars and n.s.) on the graphs show the significance comparisons to STX1A^WT^ rescue and STX1A^LEOpen^ rescue, respectively (nonparametric Kruskal-Wallis test followed by Dunn’s *post hoc* test, *p ≤ 0.05, **p ≤ 0.01, ***p ≤ 0.001, ****p ≤ 0.0001). The numerical values are summarized in Appendix Table 4.

